# From Soil to Surface: Exploring the Impact of Green Infrastructure on Microbial Communities in the Built Environment

**DOI:** 10.1101/2024.06.05.596760

**Authors:** Malin Mcgonigal, Kohei Ito

## Abstract

High microbial diversity offers extensive benefits to both the environment and human health, contributing to ecosystem stability, nutrient cycling, and pathogen suppression. In built environments, factors such as building design, human activity, and cleaning protocols influence microbial communities. This study investigates the impact of landscape design on microbial diversity and function within the "Visionary Lab" exhibition in Tokyo, Japan, using 16S rRNA gene amplicon sequencing and shallow shotgun sequencing. Despite the limited sample size, the study suggests that the Visionary Lab samples may exhibit higher microbial diversity compared to other museum areas. Potential distinct microbial community structures may be correlated with sampling locations. However, despite this, no consistent patterns were observed in virulence factors or antimicrobial resistance genes across the samples. Metabolic function analysis showed varied profiles, suggesting diverse ecological interactions influenced that may be by the curated landscape. This suggest that the curated landscape design may have the potential to enhance microbial diversity, highlighting a possible avenue to create healthier and more sustainable built environments. However, the lack of consistent patterns in virulence factors and antimicrobial resistance genes underscores the complexity of microbial community dynamics.

## Background

Recent research has increasingly recognized the importance of microbial communities in built environments, particularly their potential impact on human health and well-being. This growing field of study sits at the intersection of microbiology, architecture, and public health, offering new perspectives on how we design and manage our indoor spaces. The benefits of high microbial diversity are extensive [1, 2]. In the environment, diverse microbial communities contribute to the breakdown of organic matter and nutrient cycling, thereby maintaining ecosystem stability and productivity [3]. Soils with greater biodiversity levels are more resistant to environmental disturbances than those with reduced biodiversity, allowing ecosystems to better withstand and recover from disturbances such as pollution, climate change and habitat destruction [4]. Moreover, a complex soil microbial community increases competition for nutrients, inhibiting the development or persistence of pathogens in the soil. Thus, soil microbial diversity can directly benefit human health by suppressing disease-causing organisms, providing clean air, water, and food, and serving as a source of antibiotics [4]. In human health, it has been reported that ‘a healthy microbiota community often demonstrates high taxonomic diversity, high microbial gene richness, and stable core microbiota’ [5]. Such healthy microbiota is associated with improved immune function, reduced inflammation, improved energy and nutrient extraction, and a lower incidence of allergies and autoimmune diseases [5].

In built environments, such as homes, offices, and public spaces, microbial communities are influenced by multiple factors, including building design, ventilation systems, occupancy and human activity, types of construction materials, cleaning protocols, and external environmental conditions [6–8]. These elements interact to shape the microbial communities found in indoor spaces, affecting everything from air quality to human health. Understanding how these factors influence microbial diversity is crucial for designing healthier and more sustainable built environments [6]. The design and integration of landscape elements within built environments is one factor that can significantly influence microbial communities. Plants, in particular, have been shown to enhance microbial abundance and diversity within indoor environments [9, 10]. However, the extent to which plants influence taxonomic diversity, including genes associated with various metabolic functions or antimicrobial resistance, remains inadequately explored.

From April 2022 until August 2023, the National Museum of Emerging Science and Innovation (‘Miraikan’) in Tokyo, Japan, held an exhibition titled “Visionary Lab; Microbes Actually Are All Around”. The exhibition featured a full-scale futuristic living space with an adjoining planting area of purposefully curated floral and shrubbery arrangements. It was designed to encourage visitors to contemplate future lifestyles where humans and microbes exist plentifully in harmony. Miraikan’s planting area integrates two key concepts: multispecies sustainability and human augmentation of ecosystems. Multispecies sustainability emphasises the interconnectedness of humans, plants, and microbes, advocating for practices that support all species’ health and well-being [11]. Human augmentation of ecosystems involves deliberately enhancing of natural environments to improve ecosystem services and resilience [12].

With its landscape design, the Visionary Lab provided an opportunity to investigate the potential effects of these concepts in a real-world setting. While acknowledging the limitations of our study design, particularly the small sample size and single time point of sampling, this study aimed to provide preliminary insights into the effects of the Visionary Lab’s landscape design by comparing microbial samples from within the exhibition space to those from other areas of the Miraikan museum, using 16S rRNA gene sequencing as well as shallow shotgun sequencing to supplement our understanding of the different microbiomes. Our primary objectives were to ascertain controlled landscape design’s potential effects on the microbial ecosystem in this unique setting. We hypothesized that the specific landscape design of the Visionary Lab, particularly through the strategic integration of plant life and landscape elements, might cultivate a unique and rich microbial ecosystem that could markedly contrast with other areas within the museum. While we recognize that our study is limited in scope and cannot provide definitive conclusions, we believe it offers valuable preliminary insights that can guide future, more comprehensive research in this area. This study aims to contribute to informing urban design and public health strategies. The findings from this research may help guide future studies exploring how built environments can be designed to enhance microbial diversity, potentially contributing to improved air quality, reduced pathogen transmission, and overall public health. This approach aligns with the broader goal of sustainable urban design, which seeks to create living spaces that are resilient, ecologically balanced, and conducive to human well-being.

## Methods

### Study design, sample collection, and DNA extraction

This study employed a cross-sectional design, collecting samples from different locations within the National Museum of Emerging Science and Innovation (’Miraikan’) in Tokyo, Japan. The study included nine samples (n=9) collected from various surfaces and locations within the museum. The study was conducted at the National Museum of Emerging Science and Innovation (‘Miraikan’), located in 2--3-6 Aomi, Koto-ku, Tokyo 135-0064. Microbial samples were collected via vacuum collection or surface swabbing from different surfaces and locations, as listed in Table 1 and mapped in Figure 1. For vacuum collection, a Makita vacuum cleaner (CL107FDSH) equipped with a DUSTREAM Collector (DU-ST-1) and DUSTREAM Filter (DU-FL-1) from Indoor Biotechnologies was used. For surface swabbing, a sterile cotton-tipped swab (ESwab™; Copan Diagnostics, Brescia, Italy) was used to swab surfaces for three minutes before they were stored in tubes containing Liquid Amies Medium solution. Care was taken to ensure that collection methods did not contaminate or alter the microbial communities present. The samples were promptly sealed, stored at low temperatures, and refrigerated until DNA extraction. Extraction was performed on all nine samples using the DNeasy® PowerSoil® Pro Kit (QIAGEN, Germany) according to the manufacturer’s protocol.

**Figure 1:**
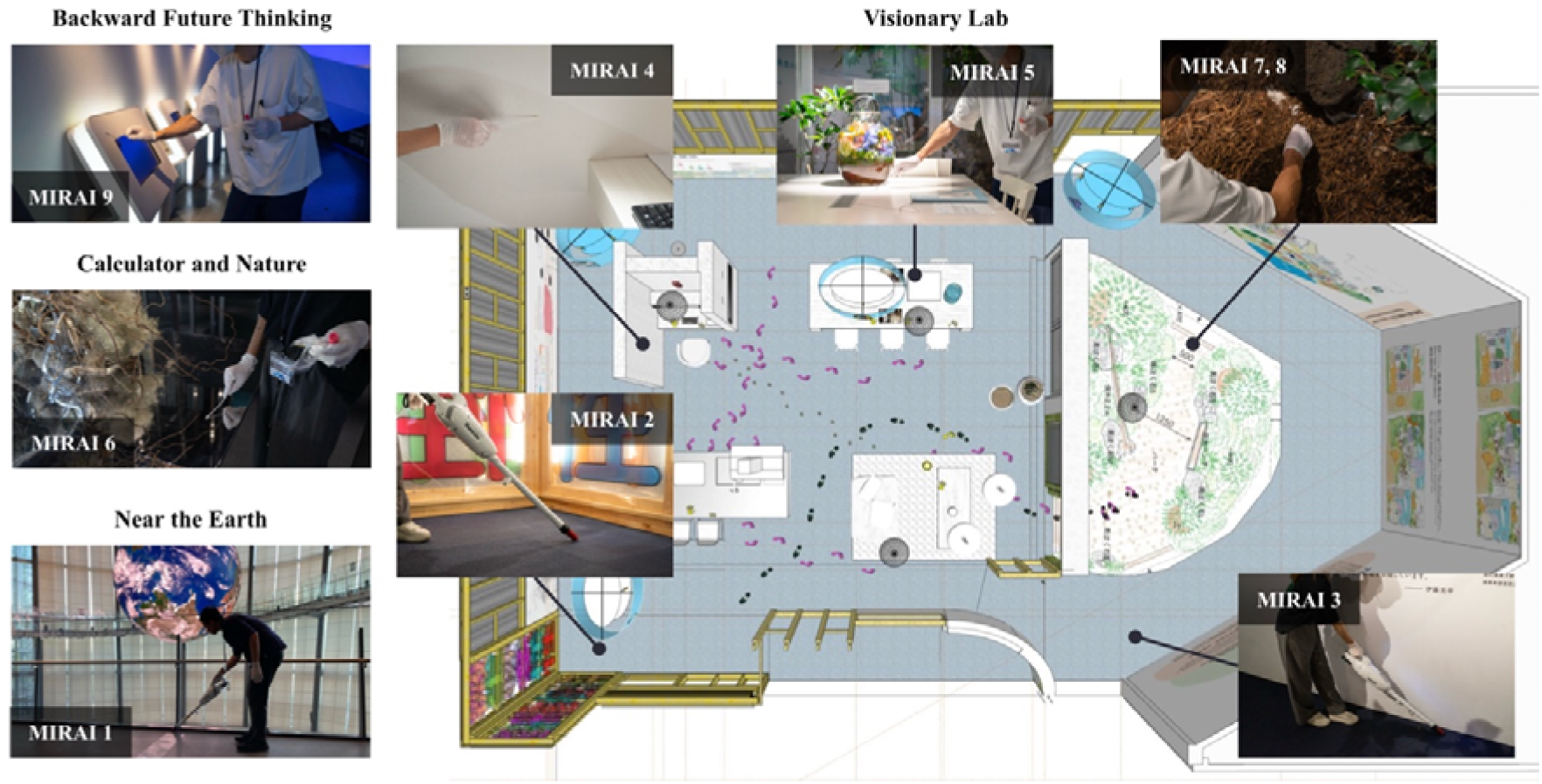
Illustration of the spatial layout of the Visionary Lab and the specific locations where microbial samples were collected. Footprints and coloured paths indicate the movement flow within the exhibit.

**Table 1:** Summary table regarding samples collected for genomic analysis, and data regarding the number of sequences processed for each sample using the two different sequencing methods. Some samples only have data for 16s rRNA amplicon sequencing as indicated by a dash in the corresponding shallow shotgun column.

| Sample | Collection protocol | Surface type | Place | Reads processed |  |
| --- | --- | --- | --- | --- | --- |
|  |  |  |  | 16s rRNA amplicon | Shallow shotgun |
| MIRAI1 | Vacuum | Floor | Near the 'Earth' | 114,111 | 21,916,203 |
| MIRAI2 | Vacuum | Floor | Visionary Lab | 107,616 | 30,083,584 |
| MIRAI3 | Vacuum | Floor | Visionary Lab near the planting area | 109,227 | 39,829,873 |
| MIRAI4 | Swab | Wall | Visionary Lab | 121,281 | - |
| MIRAI5 | Swab | Desk | Visionary Lab | 110,168 | - |
| MIRAI6 | Swab | Wall | Calculator and nature | 117,977 | - |
| MIRAI7 | Swab | Soil | Visionary Lab planting area | 104,243 | 21,768,309 |
| MIRAI8 | Swab | Soil | Visionary Lab planting area | 82,274 | 27,126,330 |
| MIRAI9 | Swab | Desk | Future backward thinking | 115,917 | - |

### Library preparation and sequencing

Sequencing libraries were generated for the 16S rRNA amplicon sequencing, and indexes were incorporated to attribute sequences to individual samples. PCR amplification of the targeted V3-V4 regions was performed using specific primers, CCTAYGGGRBGCASCAG and GGACTACNNGGGTATCTAAT. PCR products with proper size were selected via 2% agarose gel electrophoresis. An equal amount of PCR products from each sample was pooled, end-repaired, A-tailed, and then ligated with Illumina adapters. The libraries were sequenced on a paired-end Illumina platform, producing 250 base pair paired-end raw reads. Library quality was evaluated and quantified using qPCR. Quantified libraries were then pooled and sequenced on Illumina platforms according to the effective library concentration and the data amount required. DNA samples, library preparation, and amplicon sequencing were performed using 250 base paired-end sequencing on the Illumina NovaSeq 6000 system at Novogene (Beijing, China).

Sequencing libraries were prepared for the shallow shotgun metagenomic sequencing, and quality control (QC) was performed at each step to ensure data reliability. The DNA samples were randomly fragmented into shorter sequences: end-repaired, A-tailed, and ligated with Illumina adapters. These adapter-linked fragments were size-selected, PCR-amplified, and purified. The libraries were quantified using Qubit and qPCR, and their size distribution assessed using a fragment analyzer. Quantified libraries were then pooled and sequenced using the Illumina NovaSeq 6000 platform (Illumina Inc., San Diego, CA, USA) according to the effective library concentration and the required data amount, producing a 150-bp paired-end indexed recipe at Novogene (Beijing, China). Low-quality and adapter-containing reads were filtered out to ensure that only high-quality clean reads were used for downstream analysis.

### Microbiome analysis of 16S amplicon sequences

Microbiome analysis was performed based on previous studies [13, 14]. To summarise, raw FASTQ data files were imported into the QIIME2 platform (2022.8) as qza files [15] for quality control and denoising using DADA2. The sequences were organised into amplicon sequence variants (ASVs), which were then classified against the SILVA SSU 138 database using the QIIME feature-classifier classification scikit-learn package.[16, 17]. The subsequent analysis excluded ASVs classified as mitochondria, chloroplast, or unassigned. 82,274 reads were subsampled to reduce bias due to differences in read depth between samples. Shannon diversity, the number of observed features were calculated, as well as the relative species abundance. The weighted and unweighted UniFrac beta diversity indices were calculated, and the microbial community structure differences were visualised with principal coordinate analysis plots (PCoA). Data were visualised using R (version 4.0.4), ggplot2 (version 3.4.3), and ggprism (version 1.0.4).

### Microbiome analysis of shallow shotgun metagenomic sequences

Upon completion of sequencing, the Trim Galore pipeline (V0.6.1) [18] was employed for adapter trimming and quality filtering for the paired-end reads, with a quality threshold of 25 and a minimum read length of 50bp. To remove the human genome from quality-controlled sequencing, hg38 was obtained from the UCSC genome browser [19] and mapped in bowtie2 (version 2.5.4) [20]. Metagenomic assembly was carried out using Megahit (v1.2.9) [21]. Taxonomic annotations were assigned to FASTQ reads using Kraken2 (v.2.1.3) against the standard Kraken2 database, made up of NCBI taxonomic information, as well as the complete genomes in RefSeq [22]. Sequences classified as Homo sapiens were removed from the resulting taxonomy table. Metabolic functional trait profiling was performed on the assembled genomes using the METABOLIC software (v. 4.0) [23], using the METABOLIC-G implementation. Prokka (v. 1.14.5) [24] was used to annotate the prokaryotic genome, and the outputs were used by SHORTBRED (v. 0.9.4) [25] in conjunction with pre-computed shortbread markers [26] to determine the presence and absence of virulence factors and antimicrobial resistance genes across the samples against VFDB [27] and CARD [28]. Data were visualised using R (version 4.04), ggplot2 (version 3.4.2), and complexheatmap (version 3.19).

## Results

### Museum and soil environments characterised by distinct microbial communities

To understand of the taxonomic composition across different environments, we classified our samples at the genus level (16S rRNA amplicon) and at the species level (shallow shotgun). This revealed that the microbial compositions between the museum environment (MIRAI1-6, MIRAI9) and the soil environment (MIRAI7-8) were notably different, as revealed by both 16s rRNA amplicon sequencing and shallow shotgun sequencing. The amplicon sequencing results, displayed in Figure 2, suggest that the museum environment was characterized by a prevalence of genera such as *Acinetobacter* (9.73%), *Enhydrobacter* (11.31%), *Staphylococcus* (11.31%), *Bacillus* (4.41%), *Streptomyces* (1.36%), *Corynebacterium* (1.51%)*, Cutibacterium* (1.48%), and *Brevibacterium* (0.71%), while the soil environment exhibited a different microbial profile with *Pseudonocardia* (4.02%), *Streptomyces* (6.67%), *Mycobacterium* (2.81%), *Nocardioides* (1.82%), and *Steroidobacter* (0.95%) being the most prevalent. Shallow shotgun sequencing provided a more granular view, identifying species-level differences within the genera detected by amplicon sequencing (Table 2). In the museum environment, species such as *Moraxella osloensis* (8.03%), *Cutibacterium acnes* (7.81%), *Staphylococcus aureus* (3.68%), *Micrococcus luteus* (3.60%), *Brevibacterium casei* (2.616%), and *Dermacoccus nishinomiyaensis* (1.98%) were particularly abundant. Conversely, in the soil environment, species like *Pseudonocardia sp. DSM 110487* (2.14%), *Nocardioides sp. NBC_00368* (1.01%), *Streptomyces bathyalis* (0.796%), *Variovorax paradoxus* (0.70%), *Pseudonocardia sp. HH130630-07* (0.70%), and *Mycolicibacterium smegmatis* (0.68%) were observed to be dominant.

**Figure 2:**
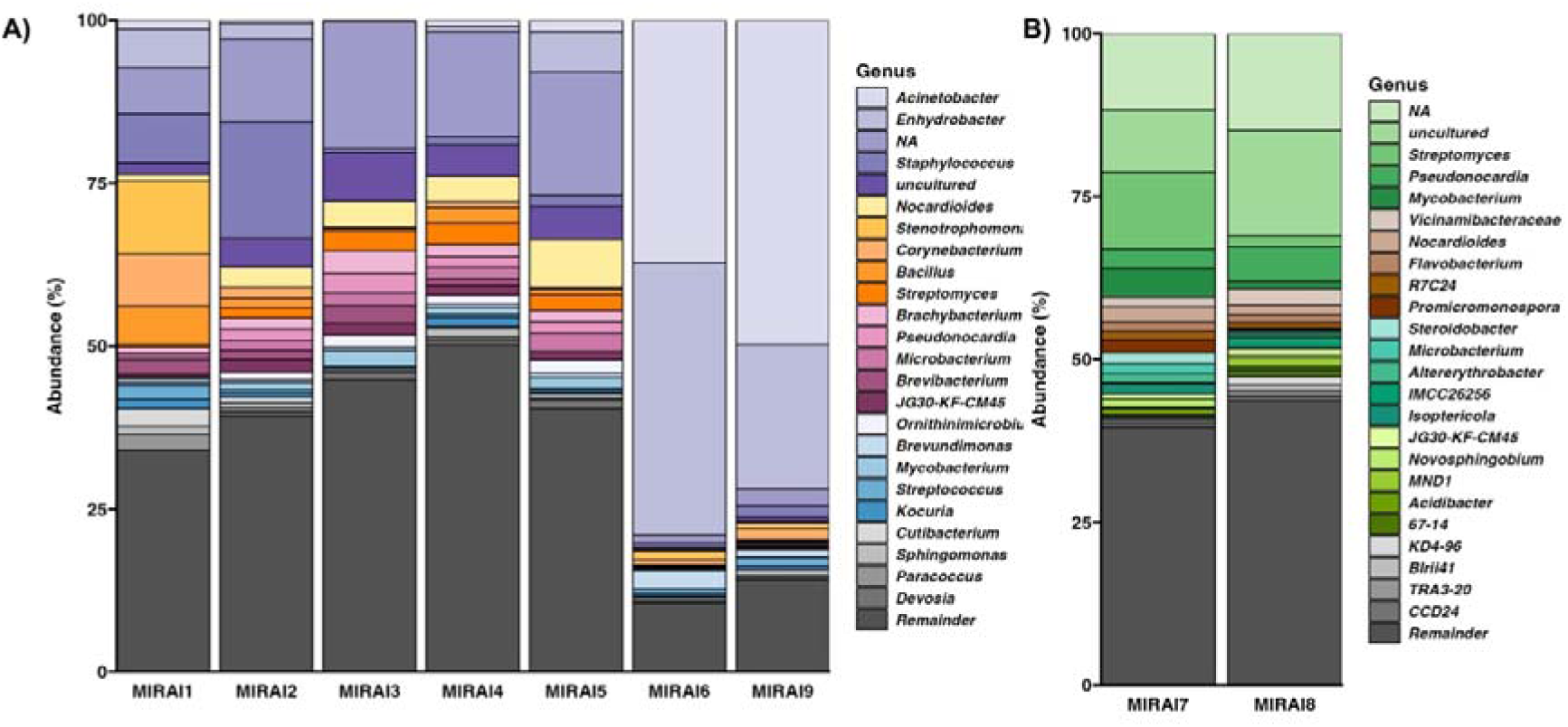
Relative abundance of most abundant microbial genera across the samples from A) the museum environment in comparison to those from B) the soil samples (within the Visionary Lab).

**Table 2:** Table displaying the relative abundance (%) of the 25 most abundant microbial species identified in samples from museum surfaces and soil samples. The ‘remainder’ row represents the cumulative relative abundance of all other species not listed in the top 25 for each sample.

| Museum microbiome |  |  |  | Soil microbiome |  |  |
| --- | --- | --- | --- | --- | --- | --- |
| 25 most abundant species | Relative abundance |  |  | 25 most abundant species | Relative abundance |  |
|  | MIRAI1 | MIRAI2 | MIRAI3 |  | MIRAI7 | MIRAI8 |
| *Moraxella osloensis* | 16.88 | 7.19 | 0.02 | *Pseudonocardia sp. DSM 110487* | 3.94 | 0.33 |
| *Cutibacterium acnes* | 14.03 | 9.35 | 0.04 | *Nocardioides sp. NBC_00368* | 1.95 | 0.07 |
| *Staphylococcus aureus* | 0.26 | 10.79 | 0.00 | *Streptomyces bathyalis* | 1.56 | 0.03 |
| *Micrococcus luteus* | 9.35 | 1.44 | 0.02 | *Variovorax paradoxus* | 0.48 | 1.02 |
| *Nocardioides sp. NBC_00368* | 0.26 | 6.47 | 2.64 | *Pseudonocardia sp. HH130630-07* | 0.15 | 1.25 |
| *Brevibacterium casei* | 7.79 | 0.00 | 0.06 | *Mycolicibacterium smegmatis* | 1.30 | 0.07 |
| *Dermacoccus nishinomiyaensis* | 5.19 | 0.72 | 0.02 | *Kribbella sp. NBC_00482* | 1.19 | 0.03 |
| *Janibacter sp. CX7* | 0.78 | 4.32 | 0.50 | *Pseudonocardia sp. AL041005-10* | 0.15 | 1.05 |
| *Corynebacterium tuberculostearicum* | 4.68 | 0.72 | 0.00 | *Pseudoxanthomonas suwonensis* | 0.93 | 0.26 |
| *Janibacter limosus* | 0.52 | 4.32 | 0.32 | *Stackebrandtia nassauensis* | 1.10 | 0.03 |
| *Staphylococcus hominis* | 4.16 | 0.72 | 0.00 | *Mycolicibacterium moriokaense* | 1.02 | 0.07 |
| *Agrococcus carbonis* | 0.26 | 4.32 | 0.11 | *Streptomyces xanthii* | 1.04 | 0.03 |
| *Sphingomonas sp. FARSPH* | 3.12 | 1.44 | 0.06 | *Ramlibacter tataouinensis* | 0.13 | 0.89 |
| *Janibacter terrae* | 0.52 | 3.60 | 0.26 | *Enhydrobacter sp.* | 0.26 | 0.69 |
| *Pseudonocardia sp.*<br>*DSM 110487* | 0.00 | 1.44 | 2.26 | *Pseudonocardia sp.*<br>*HH130629-09* | 0.11 | 0.79 |
| *Brachybacterium sp.*<br>*SGAir0954* | 0.26 | 2.16 | 1.18 | *Sorangium cellulosum* | 0.37 | 0.46 |
| *Janibacter melonis* | 0.52 | 2.16 | 0.88 | *Streptomyces nigra* | 0.78 | 0.03 |
| *Nocardioides sp.*<br>*JQ2195* | 0.00 | 2.16 | 1.38 | *Usitatibacter rugosus* | 0.09 | 0.72 |
| *Corynebacterium*<br>*afermentans* | 3.38 | 0.00 | 0.00 | *Variovorax sp. HW608* | 0.35 | 0.46 |
| *Dermacoccus sp.*<br>*PAMC28757* | 2.60 | 0.72 | 0.02 | *Luteitalea sp. TBR-22* | 0.17 | 0.62 |
| *Janibacter sp. AIS7* | 0.26 | 2.88 | 0.07 | *Croceibacterium sp.*<br>*D39* | 0.37 | 0.43 |
| *Brachybacterium sp.*<br>*NBEC-018* | 0.00 | 2.16 | 0.97 | *Lysobacter soli* | 0.56 | 0.23 |
| *Janibacter indicus* | 1.30 | 1.44 | 0.19 | *Luteitalea pratensis* | 0.15 | 0.56 |
| *Mycolicibacterium*<br>*smegmatis* | 0.00 | 0.72 | 1.79 | *Kribbella sp.*<br>*NBC_00709* | 0.67 | 0.03 |
| *Nocardioides sp. zg-*<br>*1228* | 0.00 | 1.44 | 0.95 | *Lacunisphaera*<br>*limnophila* | 0.26 | 0.43 |
| Remainder | 23.90 | 27.34 | 86.26 | Remainder | 80.92 | 89.43 |

### Planting areas contributed to increasing microbial diversity within the exhibition

In order to generate further insights into the effectiveness of landscape design on microbial diversity, we analysed the alpha diversity metrics. The alpha diversity indices, as shown in Table 3, indicate notable differences between the Visionary Lab samples (MIRAI2, MIRAI3, MIRAI4, MIRAI5, MIRAI7, and MIRAI8) and the non-Visionary-Lab samples (MIRAI1, MIRAI6, and MIRAI9). The Shannon diversity indices for the Visionary Lab samples tended to be higher, ranging from 9.28 to 10.48, compared to the non-Visionary-Lab samples, which ranged from 4.87 to 7.89. This suggests a potentially greater diversity of microbial species in the Visionary Lab environment. Similarly, the number of observed features was markedly higher in the Visionary Lab samples, with counts between 1,556 and 2,217, while the non-Visionary-Lab samples showed lower counts ranging from 384 to 1,092. The phylogenetic diversity (Faith PD) also appeared to reflect this trend, with the Visionary Lab samples exhibiting values from 85.99 to 150.13, higher than the non-Visionary-Lab samples, which ranged from 38.27 to 69.42. Additionally, the Pielou evenness values are generally higher for the Visionary Lab samples, mostly above 0.87, potentially indicating a more even distribution of species within these microbial communities, whereas the non-Visionary-Lab samples displayed lower evenness values ranging from 0.56 to 0.78. Within all samples and across all metrics, MIRAI8 (a soil sample from the visionary lab) shows the highest overall diversity.

**Table 3:** Alpha diversity metrics for all samples. Metrics include Shannon diversity (measure of species richness and evenness), observed features (number of distinct features observed), Faith’s Phylogenetic Diversity (measure of biodiversity that incorporates phylogenetic differences between species), and Pielou’s Evenness (measure of how evenly the species are distributed).

| Alpha diversity metrics |  |  |  |  |
| --- | --- | --- | --- | --- |
| Sample | Shannon diversity | Observed features | Faith PD | Pielou evenness |
| MIRAI1 | 7.893 | 1,092 | 69.428 | 0.782 |
| MIRAI2 | 9.281 | 1,597 | 112.933 | 0.872 |
| MIRAI3 | 9.736 | 1,694 | 105.363 | 0.908 |
| MIRAI4 | 9.955 | 1,616 | 88.141 | 0.934 |
| MIRAI5 | 9.368 | 1,556 | 85.999 | 0.883 |
| MIRAI6 | 4.877 | 384 | 38.279 | 0.568 |
| MIRAI7 | 9.991 | 1,702 | 86.597 | 0.568 |
| MIRAI8 | 10.488 | 2,217 | 150.135 | 0.944 |
| MIRAI9 | 5.702 | 524 | 44.321 | 0.631 |

### Location-Based Microbial Clustering in Museum Samples

To further understand the differences in microbial community structures, we performed beta diversity analysis using weighted UniFrac Principal Coordinate Analysis on samples analysed by 16s rRNA amplicon sequencing. The results, displayed in Figure 3, reveal clear clustering of microbial communities that correspond to their specific sampling locations within the museum, as mapped in Figure 1.

**Figure 3:**
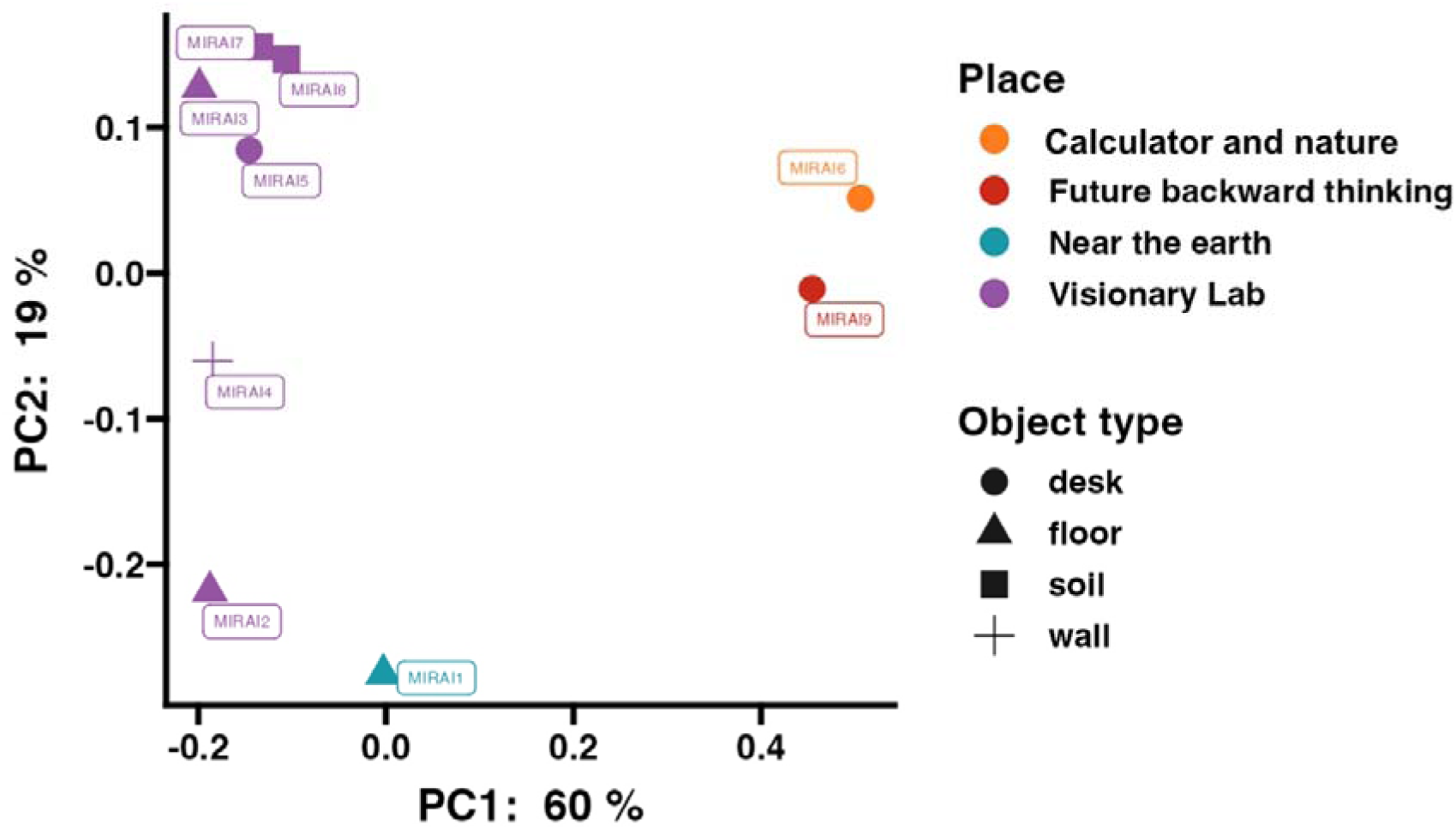
Weighted unifrac Principal Coordinate Analysis (PCoA) for samples analysed by 16S rRNA amplicon sequencing. Samples are colour-coded by location and shaped according to surface type.

The PCoA plot can be broadly separated into three main clusters. The first cluster includes samples MIRAI6 and 9, taken from the “Future Backward Thinking” and “Calculator and Nature” exhibits, respectively. These exhibits are located close to each other, as shown on the map, which may explain to their similar microbial community structures. The second cluster consists of samples MIRAI2 and MIRAI1, which again are located close to one another (whilst MIRAI2 was taken from within the Visionary Lab exhibition, it was towards the entrance/exit). The third cluster, comprising the remaining samples, were all taken from various locations within the main area of the visionary lab exhibit, and this proximity is reflected in their similar microbial community compositions.

### No consistent patterns observed in the presence of virulence factors or antimicrobial resistance genes

Having understood the structure of the microbiome of our samples; in order to gain further insights into the implications, we performed analysis of virulence factors and antimicrobial resistance. Table 4 presents a comparative analysis of virulence factors and antimicrobial resistance between samples taken from within the Visionary Lab (MIRAI2-8), where landscape design was implemented, and an external sample (MIRAI1). Sample MIRAI1, collected outside the premises, exhibited a moderate presence of virulence factors with notable counts in elements such as fumarate hydratase (VFDBID: VFG038025) and transposase (virulence|961). However, no antibiotic resistance genes were detected in this sample. Among the Visionary Lab samples, sample MIRAI3 showed a significant presence of multiple virulence factors with higher counts and hits, such as biopolymer transporter ExbD (gi 406029600), an uncharacterised protein previously found in *Penicillium rubens* (gi 15839882) and M28 family peptidase (gi 120401443). Sample M3 also contained antibiotic resistance genes *RbpA* and *Mycobacterium*. Interestingly, sample M7, collected from the soil around the planting area, showed a presence of virulence factors like phosphoribosyl-AMP cyclohydrolase (VFDB ID: VFG028663), UV excision repair protein RAD23 homolog B isoform X1 (VFDB ID: VFG009731), and the same uncharacterised protein identified in MIRAI3 (gi 15839882). It also shared the same antibiotic resistance gene as MIRAI3-*RbpA* (ADV91011). Sample MIRAI8, also from the soil around the planting area, shows no detectable virulence factors or antibiotic resistance genes.

**Table 4:**
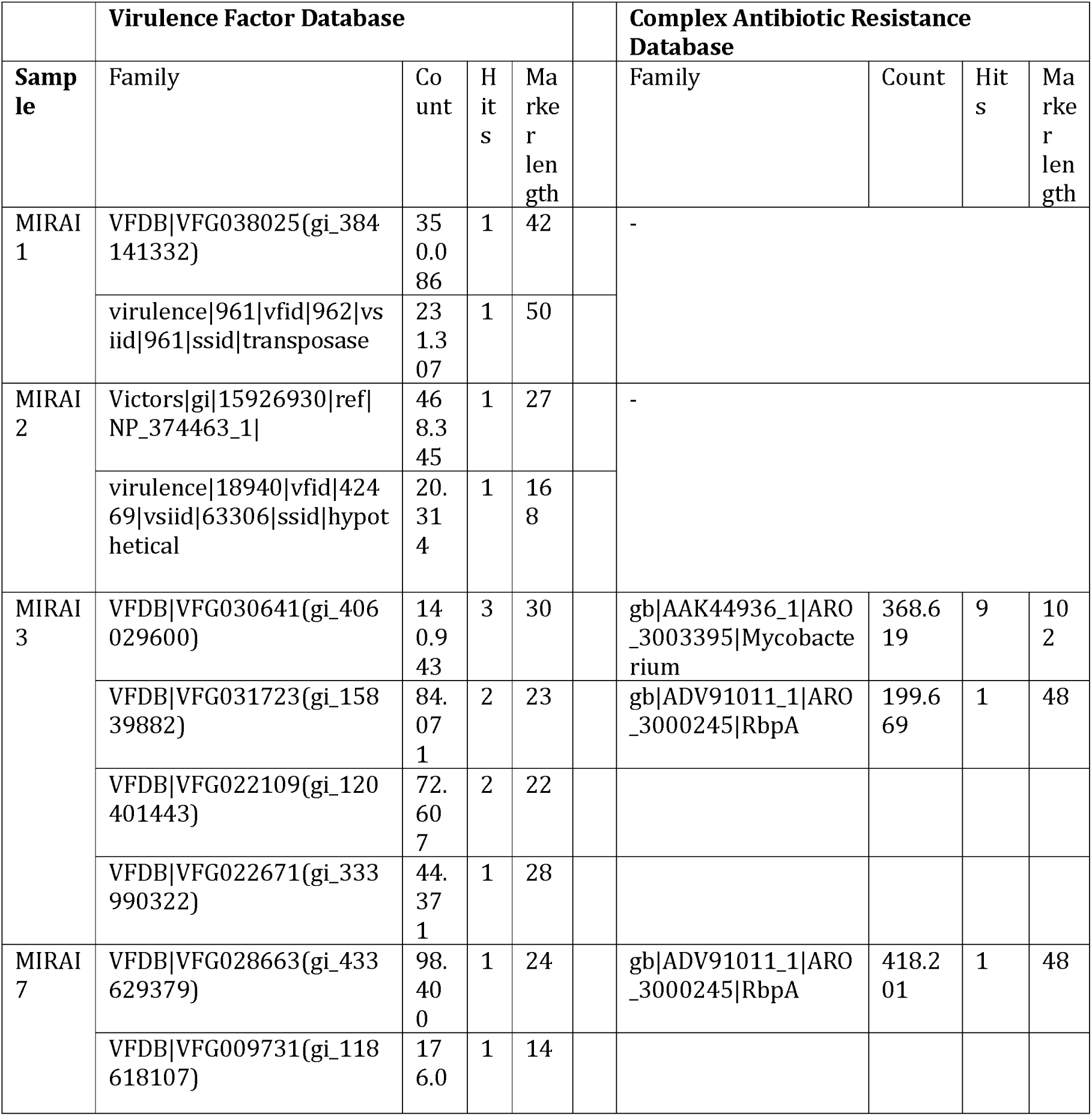
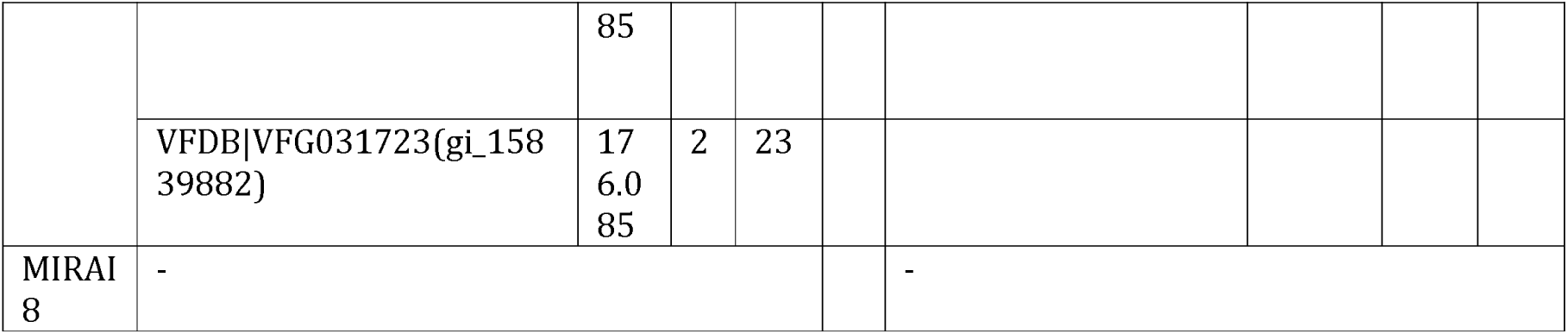
Virulence factors and antibiotic resistance genes identified in samples sequenced by shallow shotgun sequencing. ‘Family’ represents the database family and gene identifier, ‘count’ is the total abundance count of each gene, ‘hits’ is the number of times the gene was detected, and ‘marker length’ is the length of the gene marker.

### Planting areas contribute to microbiomes with a larger array of metabolic functions

Having not been able to elucidate any concrete patterns in virulence or antimicrobial resistance, we next strived to further understand the functional landscape of the microbiomes. The heatmap in Figure 4 illustrates the presence or absence of genes coding for various metabolic functions across different samples using a shallow shotgun sequencing approach. Overall, the samples exhibited some common patterns but also displayed distinct differences. Categories like "Metal reduction" and "Chlorite reduction" appeared consistently across all samples, indicating these metabolic functions are widely shared. However, other categories, such as "Complex carbon degradation" and "Nitrogen cycling," show significant variation between samples, suggesting differing metabolic capabilities. Sample MIRAI3, with a high presence of blue, indicates many metabolic functions, suggesting it might be the most diverse in terms of metabolic activity. Conversely, samples MIRAI1 and MIRAI2 have the least amount of blue, indicating fewer metabolic functions and suggesting limited metabolic diversity. Samples MIRAI7 and MIRAI8, collected from the soil around the planting area, showed moderate metabolic diversity, with MIRAI7 exhibiting a broad range of metabolic capabilities, including complex carbon degradation and nitrogen cycling, pointing to a rich microbial community.

**Figure 4:**
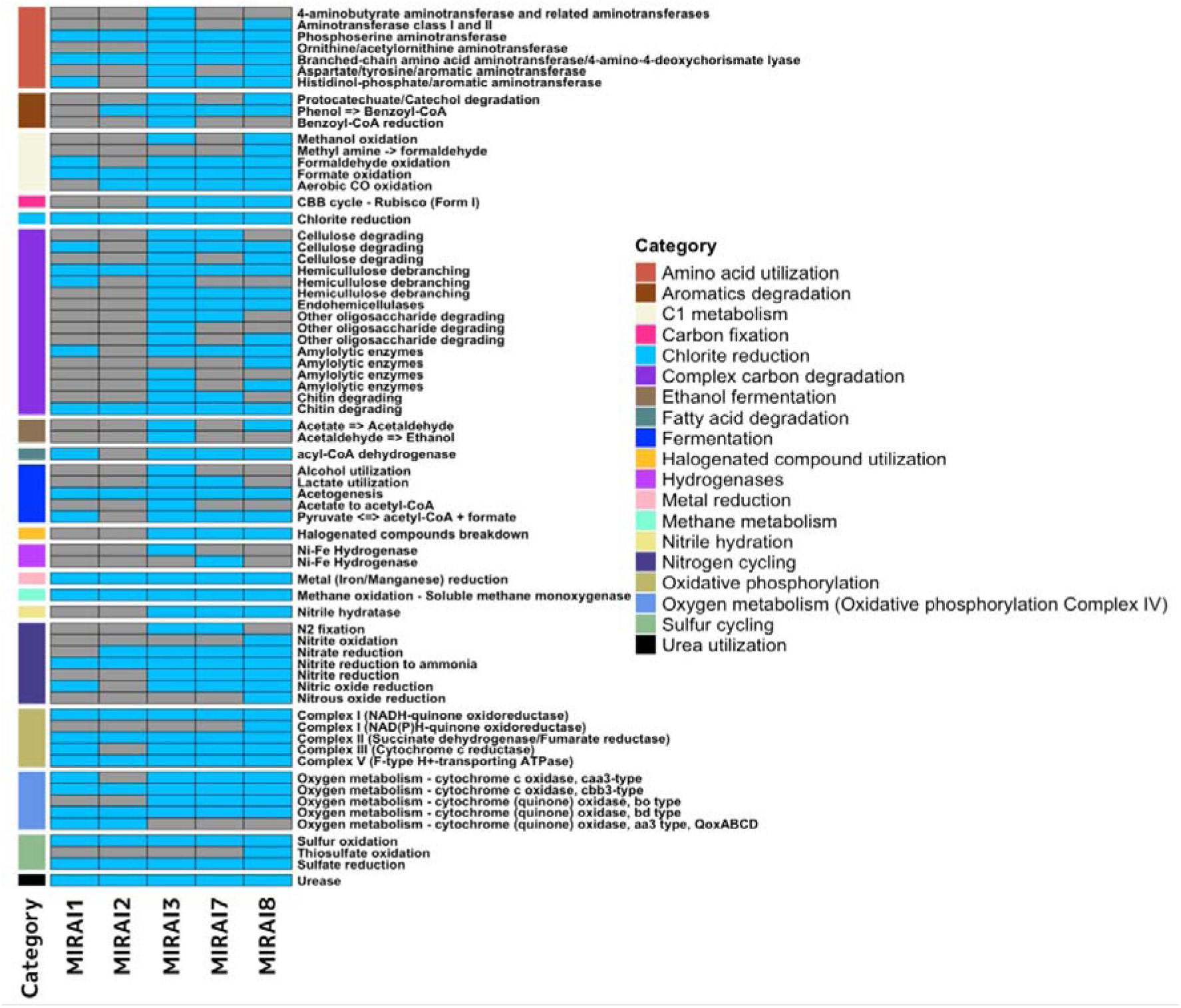
Heatmap illustrating the presence or absence of genes coding for various metabolic functions across samples sequenced by a shallow shotgun approach. The legend on the right categorises each function by its metabolic category, aiding in the interpretation of metabolic diversity and specialisation among the samples.

## Discussion

### Microbial composition and diversity

The observed distinct microbial profiles between the museum and soil samples can be attributed to the differential ecological niches and associated activities within these settings. These profiles appear to reflect the influence of different environmental contexts. The museum environment, characterised by genera such as *Acinetobacter*, *Enhydrobacter*, *Staphylococcus*, *Bacillus*, *Streptomyces*, *Corynebacterium*, *Cutibacterium*, and *Brevibacterium*, predominantly appears to harbour human-associated microbes (Figure 2). These taxa are commonly found on human skin, surfaces frequently touched by humans, and in indoor air, potentially reflecting the high level of human activity and interaction within the museum environment [29]. For example, *Cutibacterium acnes*, which was notably abundant in the museum samples, is a well-known skin commensal-the most prevalent bacteria residing in human, healthy, sebum-rich skin areas such as the face and back [30]. Similarly, *Staphylococcus aureus and Staphylococcus hominis*, are also human-associated bacteria that thrive in indoor environments, and their presence in the museum samples aligns with previous studies reporting that *staphylococci*, usual colonisers of the human skin and upper respiratory tract, are commonly spread from humans to built environment surface and air [31–33]. *Moraxella osloensis* was found to be the most abundant species across the exhibition samples, and is a gram-negative aerobic bacterium that is considered to be an opportunistic human-associated pathogen, previously isolated from the upper respiratory tract, blood, genitourethral speciments, and other sites in humans [34–36]. Again, *Moraxella osloensis* has been found in several locations within the indoor environment where humans frequent, including sinks and laundry rooms [35, 37]. *Corynebacterium* is another predominant genus in the skin microbiome, with *Corynebacterium tuberculostearicum* being a ubiquitous skin-colonising bacterium which has been found to play a role in skin health and disease [38, 39]. These findings are consistent with previous studies that have shown a high prevalence of human-associated microbes in indoor environments with significant human traffic [29, 40, 41].

In contrast, the soil samples from the Visionary Lab’s planting area exhibited a different microbial profile characterized by environment-associated genera such as *Pseudonocardia, Streptomyces, Mycobacterium, Nocardioides*, and *Steroidobacter* (Figure 2). These taxa are typically associated with soil and plant materials, indicating the influence of natural environmental conditions in shaping microbial community composition. For example, members of the genus *Pseudonocardia* are commonly found in soil, plants and the wider environment [42]. In particular, *Pseudonocardia sp.* DSM 110487 and *Pseudonocardia sp. HH 130630-07,* two species identified by shallow shotgun sequencing to be abundant in the soil samples, are known for their capabilities in organic matter degradation and their symbiotic relationships with insects, respectively, contributing to the soil’s health and ecological balance [43, 44]. Regarding the genera *Streptomyces,* of which *Streptomyces bathyalis* was identified as an abundant species within our soil microbiome, studies support our findings, showing that that the bacteria belonging to this genus are mainly found in soil but are also occasionally isolated from manure and other environmental sources [45]. The presence of *Mycobacterium* and *Nocardioides* further supports the environmental origin of our samples, as species from these genera are again frequently isolated from soil and are involved in nutrient cycling and biodegradation processes [46, 47]. By understanding the distinct microbial compositions in different contexts, this study begins to shed light on the importance of considering both human and environmental factors in designing and managing indoor spaces to enhance microbial diversity and promote ecological balance.

The small discrepancies observed between the results obtained from 16s rRNA amplicon sequencing and shallow shotgun sequencing regarding the most abundant genera and species can likely be attributed to the limitations and biases inherent in each method. While amplicon sequencing provided a broad overview of the dominant genera, it was subject to PCR biases that can skew the representation of microbial communities [48]. Additionally, the lack of comprehensive reference databases for soil microbes likely resulted in the underrepresentation of many species under both sequencing methods [49]. Whilst shallow shotgun sequencing offered higher resolution at the species level, uncovering specific species within the detected genera, this method may also have been subjected to potential biases in DNA extraction and library preparation steps. These findings underscore the importance of using complementary sequencing methods to achieve a comprehensive characterization of microbial communities. By integrating data from both 16S rRNA amplicon and shallow shotgun sequencing, we can obtain a more accurate and nuanced picture of the microbial ecosystems present in different environments [50, 51]. Furthermore, understanding the distinct microbial profiles associated with human-dominated indoor settings versus natural soil environments can inform strategies for managing microbial diversity and maintaining healthy indoor microbiomes.

Our findings suggest a potential impact of landscape design on microbial diversity within the Visionary Lab compared to other areas of the Miraikan museum. While limited by our small sample size, consistently suggest higher microbial diversity in the Visionary Lab samples compared to the non-Visionary-Lab samples (Table 3). Higher Shannon diversity indices in the Visionary Lab compared to the non-Visionary-Lab suggest a more complex microbial community [52]. This higher diversity is further supported by the increased number of observed features and higher phylogenetic diversity values, indicating a broader range of microbial species and greater evolutionary variety in the Visionary Lab, which could potentially enhance ecosystem multifunctionality [53]. The higher Pielou evenness values in the Visionary Lab samples indicate a more balanced distribution of species, suggesting that the environment supports a wide range of microbial taxa without dominance by a few species [54]. Higher evenness within microbial communities is often linked to enhanced ecosystem stability and productivity [55]. This evenness may be attributed to the diverse plant life and landscape elements designed to promote a harmonious coexistence of microbes. The notably high diversity observed in sample MIRAI8, a soil sample from the Visionary Lab, further supports our hypothesis about the effectiveness of the landscape design in enhancing microbial richness and evenness. This finding aligns with supports existing literature which consistently suggests that soil, a critical component of landscape design, plays a crucial role in supporting a diverse microbial ecosystem [56]. Similarly, a study by van der Heijden et al. (2008) found that plant diversity enhances ecosystem productivity and stability, partly through its effects on soil microbial communities [57].

The samples outside the Visionary Lab exhibited lower microbial diversity, as evidenced by their alpha diversity indices (Table 3). Several factors may contribute to this. Firstly, the lack of intentional landscape design outside the exhibition means there are fewer plants and less environmental complexity to support a diverse microbial community. As discussed, studies have shown that plant diversity and structural complexity provide a range of niches and resources, which enhance microbial diversity. Without these elements, it is expected that microbial communities will be less diverse. Secondly, human activity in these areas might introduce human-associated microbes while reducing environmental microbes -something we can confirm through our taxonomic assessment (Figure 2). These high-traffic areas can experience frequent cleaning and disinfection, which can diminish microbial diversity by selectively eliminating certain taxa [58]. Additionally, our taxonomic assessment revealed that there were few human-associated microbes which appeared to dominate in the non-Visionary-Lab environments, leading to lower overall diversity (reflected in the observed features metric). Thirdly, the environmental conditions outside the Visionary Lab might not be as conducive to microbial diversity. Factors such as lower humidity, less organic matter, and fewer plant-derived resources can limit microbial growth and diversity [59].

The beta diversity analysis provides significant insights into the spatial organisation and transfer mechanisms of microbial communities within the museum. The observed clustering of microbial communities, corresponding to their sampling locations, suggests the influence of spatial proximity on microbial diversity (Figure 1 and Figure 3). These findings highlight the possible importance of environmental features and the physical layout in shaping microbial ecosystems. Mechanisms of microbial transfer may play a crucial role in explaining the formation of these distinct clusters. Human activity is a primary driver of microbial dispersal within built environments [6]. As visitors move through the museum, they inadvertently carry and deposit microbes through skin, clothing, and respiratory emissions, contributing to microbial transfer [7] This human-mediated transfer is particularly evident in areas with high foot traffic, such as the entrances and exits of exhibits. For example, the clustering of samples MIRAI2 and MIRAI1 could potentially be attributed to the high visitor traffic near the Visionary Lab entrance, facilitating the homogenization of microbial communities in these regions (Figures 1 & 3). Surface contact represents another critical mechanism [6]. Surfaces such as exhibit materials, floors, and interactive displays serve as reservoirs for microbial communities. Visitors touching these surfaces enhance microbial exchange, particularly in areas with interactive exhibits. Human activity is a significant vector for microbial transfer, as people can carry microbes on their skin, clothing, and personal items [6]. The interaction between visitors and the plant-rich environment of the Visionary Lab likely facilitates the transfer and dissemination of diverse microbial species. Previous studies have shown that environmental conditions, such as moisture levels, nutrient availability, and vegetation cover, can significantly impact microbial diversity and composition [60, 61]. The clustering of samples within specific exhibits suggests that the unique conditions and interactions within these microenvironments drive distinct microbial assemblages. Although in this given study we did not collect data on airflow or ventilation, it is known that airborne dispersal can also contribute to microbial transfer from different surfaces -for example transfer from soil samples to non-soil samples [62]. Soil and plant surfaces themselves are reservoirs of microbial life, and their interaction with the surrounding environment can lead to the spread of microbes, where air currents and ventilation systems lift microbes from soil into the area and deposit them onto nearby surfaces [63]. This could be the case in the Visionary Lab, where the strategic placement of environmental features, such as plants, create microhabitats that support diverse microbial populations, contributing to the unique microbial ecosystems observed within the Visionary Lab samples.

The clustering patterns observed in the beta diversity analysis are consistent with the theory that proximity to curated environmental features significantly influences microbial composition [64]. This correlation reinforces the idea that strategic environmental design can be leveraged to shape microbial ecosystems in desired ways. By understanding the mechanisms of microbial transfer, such as human activity, airflow, and surface contact, we can optimise the design of built environments to promote beneficial microbial communities. In conclusion, the distinct community structures revealed by the beta diversity analysis highlight the importance of spatial organisation and environmental design in shaping microbial ecosystems within the museum.

Despite the valuable insights gained, it must be acknowledged that due to the small sample size (n=9), of which only 5 were sequenced by shallow shotgun methods, it may not have been possible to fully capture the variability and complexity of microbial communities within the Visionary Lab and the Miraikan museum. A larger sample size would provide a more comprehensive understanding of the microbial diversity in these environments and allow for further statistical tests to validate the significance of findings.

### Virulence factors, antimicrobial resistance, and metabolic function profiles

The comparative analysis of virulence factors and antimicrobial resistance between samples revealed no clear consistent patterns, highlighting the complexity of microbial community dynamics on human health (Table 4). The prevalence of virulence factors in sample MIRAI1 suggests a baseline level of microbial pathogenicity in the external environment, whilst the absence of antibiotic resistance genes in this sample indicates a potentially lower risk of antibiotic-resistant infections in these settings. The significant presence of virulence factors in MIRAI3 suggests that microbes in this controlled environment have adapted mechanisms to enhance their survival and pathogenicity. ExbD, for instance, is involved in nutrient uptake and energy transduction, which could enhance microbial fitness in nutrient variable conditions. It has been identified before from *Verrucomicrobiota* bacterium, which in turn has been commonly isolated from environmental environments such as marine and soil locations, as well as the human microbiome such as human faeces [65]. This makes sense in the context of our results, given it was found in sample MIRAI3 which is a sample taken from a surface-so lots of contact with human environments, but close to the planting area-so transfer from the soil samples. The detection of antibiotic resistance genes, such as *RbpA* and *Mycobacterium* in this sample also raises concerns about the potential for these environments to harbour and disseminate resistant microbes. *RbpA* is known to provide resistance to rifampicin, a critical antibiotic for treating tuberculosis [66]. This gene indicates that even in curated environments, there is a risk of developing and spreading antimicrobial resistance, likely due to selective pressures and microbial interactions. MIRAI 7, collected close to MIRAI 3 from the soil around the planting area, shared several virulence factors and the RbpA resistance gene with MIRAI3. This suggests that the landscape design, which includes soil and plant materials, creates microhabitats that support diverse microbial communities with both pathogenic and resistant strains. Phosphoribosyl-AMP cyclohydrolase, detected in MIRAI7, is crucial for nucleotide biosynthesis and microbial proliferation, indicating that soil microbes are well-equipped for growth and survival in these enriched environments [67]. The absence of detectable virulence factors and antibiotic resistance genes in MIRAI8 could be due to localised differences in soil composition or microbial interactions that limit the growth of pathogenic and resistant strains.

A small number of studies [68–71] have concentrated on public settings in built-up environments, but the majority of these studies have demonstrated that AMR resistance is highly abundant in public settings across various species [72]. One study revealed that highly maintained environments (subject to intensive cleaning and disinfection processes) exhibit a higher diversity of antimicrobial resistance genes and reduced microbial diversity [73]. Based on this, we hypothesised that the soil samples taken from the planting area within the visionary lab would show the least AMR genes and virulence factors (due to its high microbial diversity) and those with more human contact (samples taken from surfaces) would have more AMR genes and virulence factors. However, overall, these results suggest that the influence of landscape design on virulence factors and antimicrobial resistance is complex and does not follow a clear, consistent pattern. This may be due to our study’s limited sample size and scope, which does not fully capture the broader patterns and interactions within the microbial communities. PROKKA and SHORTBRED are tools which depend on the quality and completeness of the reference databases and the contigs used in the analysis. Incomplete or fragmented contigs may lead to underestimation or misidentification of genes, affecting the results. The variability observed emphasised the need for further research to understand the multifaceted interactions between environmental conditions, microbial communities, and human activity in shaping the presence and distribution of virulence factors and antimicrobial resistance genes in built environments.

The distinct metabolic profiles highlight how controlled landscape design can shape microbial communities, fostering unique ecological interactions and functions (Figure 4). Samples which share genes that code for common functions can be attributed to these functions’ broad applications in various environmental and human settings. Within the group of functions which exhibit notable variation within the samples (function present in 30-70% of samples), the most common pattern is for the function to be present in MIRAI3, MIRAI7, and MIRAI8 and not present within MIRAI1 and MIRAI2. For example, the complex carbon degradation category follows this pattern. This distribution pattern of these genes across the samples can be explained by the functions within the category being central in an environment with plant material (e.g., cellulose-degrading, hemicellulose-debranching, etc…) [74]. As a result, genes which code for these functions are more likely to be found within the soil/plant samples (MIRAI7 and MIRAI8). The reason these genes are also found to be present in sample MIRAI3, could be attributed to microbial transfer over short distances-as MIRAI3 was taken from a location close to the planting area where MIRAI7 and MIRAI8 were taken. This same explanation can be applied to virtually all the remaining functions within this variation category within samples. Fermentation is another function that demonstrates significant variation across samples. Genes associated with fermentation are predominantly found in plant/soil samples (MIRAI7 and MIRAI8) and are largely absent in samples from human-associated environments (MIRAI1 and MIRAI2).

This analysis focuses solely on the presence or absence of genes coding for specific functions, which presents certain limitations. For example, in the context of ’nitrogen cycling’, the pattern of presence or absence of genes varies across the five different functions, leading to the question of why one sample contains one type of gene while lacking genes for closely related functions in the same nitrogen cycle. These ambiguities may arise from biological factors or contamination, complicating interpretation. Thus, more comprehensive analyses considering specific gene abundance are needed better to understand the genetic landscape and its functional implications. Nonetheless, the broad application of certain metabolic functions across various settings underscores their ecological importance, while other variability points to localised environmental influences.

## Conclusion

This study comprehensively outlines the impact of green infrastructure on microbial diversity in the context of the Visionary Lab compared to other areas of the Miraikan museum. The curated landscape design, incorporating diverse plant life, supports a more complex and balanced microbial community, contributing to enhanced microbial richness, evenness, and ecosystem multifunctionality. While the Visionary Lab samples exhibited higher microbial diversity, the study did not reveal consistent patterns in virulence factors or antimicrobial resistance genes. This variability underscores the complexity of microbial community dynamics and the influence of localised environmental conditions. Future studies should increase the sample size and variety of sampling locations within and outside the Visionary Lab and employ further advanced sequencing techniques (such as deep metagenomic sequencing) to enhance the robustness and generalizability of results. Additionally, exploring the impact of different environmental variables, such as humidity, temperature, and light exposure, on microbial communities, and conducting longitudinal studies to track microbial changes over time would further elucidate the dynamics of microbial ecosystems. Using this comprehensive approach to build upon the findings from this study would contribute to developing evidence-based guidelines for creating microbial-friendly and health-promoting indoor environments, aligning with the broader goal of sustainable urban design.

## Declarations

Ethics approval and consent to participate:

Not Applicable.

## Consent for publication

Not Applicable

## Authors’ contributions

K.I. designed the study and analysed the 16s rRNA Amplicon data. M.M. interpreted the 16s rRNA Amplicon data and analysed and interpreted the shallow shotgun data. M.M was the major contributor in writing the manuscript. All authors read and approved the final manuscript.

## Acknowledgements

We would like to extend our sincere gratitude to Marin Yamaguchi and Keisuke Otsuka for their diligent work in collecting the samples. Our thanks go to the National Museum of Emerging Science and Innovation (Miraikan) for hosting the "Visionary Lab; Microbes Actually Are All Around" exhibition between April 2022 and August 2023 and for allowing us to collect samples within their facility (https://www.miraikan.jst.go.jp/resources/visionarieslab/visionaries-archive-003.html). We also thank the Japan Science and Technology Agency (JST) for their collaboration and support in the Visionary Lab exhibition. This study would not have been possible without their collaboration and support.

## Availability of Data and Materials

The datasets generated through 16S rRNA amplicon and metagenomic sequencing are available and deposited in the DDBJ Sequence Read Archive (DRA) database under accession numbers DRR582054-DRR582069 and BioProject PRJDB18223.

